# Neutrophils Restrain CCR2-Dependent Monocyte Immunopathology During Pulmonary Cryptococcosis

**DOI:** 10.64898/2026.10.26.762039

**Authors:** Mikayla C. Harden-Pruitt, Rebekah G. Watson, Camaron R. Hole

## Abstract

Neutrophils are essential for host defense against most invasive fungal pathogens, yet their role in cryptococcosis remains paradoxical. Neutropenia is not a recognized risk factor for cryptococcal disease, and an early study reported that neutrophil depletion improves survival in mice, supporting the prevailing view that these cells are dispensable. Here we re-examined this question using Ly6G-specific depletion strategies. Contrary to prior reports, neutrophil-depleted mice infected with *Cryptococcus neoformans* succumbed significantly more rapidly than isotype-treated controls. Accelerated mortality occurred without any increase in pulmonary fungal burden at 24h, 48h, or day 7, or in brain and spleen at day 7, indicating host-mediated rather than pathogen-driven death. This phenotype was independent of mouse genetic background, depletion antibody, inoculum, and fungal strain. Within 24h, neutrophil depletion provoked an early chemokine and IL-17A response, followed by pronounced pulmonary accumulation of inflammatory monocytes and macrophages by day 7. Strikingly, the increased mortality associated with neutrophil depletion was completely absent in *Ccr2*^−/−^ mice, implicating CCR2-dependent inflammatory responses in disease severity. Together, these findings reveal a previously underappreciated protective function of neutrophils during cryptococcosis, whereby they constrain inflammatory monocyte-associated immunopathology and promote host survival independently of measurable changes in fungal burden.

**IMPORTANCE:** *Cryptococcus neoformans* causes life-threatening infections, particularly in individuals with weakened immune systems. Neutrophils are white blood cells that protect against many fungal infections, but their role in cryptococcosis has remained unclear. We found that depleting neutrophils from infected mice consistently accelerated mortality without increasing fungal burden in their tissues. Instead, neutrophil depletion triggered excessive lung inflammation and accumulation of inflammatory monocytes. Mice that could not recruit inflammatory monocytes into the lungs were completely protected, identifying those cells as the cause of lethal damage. Neutrophils therefore act as a brake on a damaging inflammatory response, highlighting the potential of targeting host immune responses to improve outcomes during cryptococcosis.

## INTRODUCTION

Cryptococcosis remains a major cause of morbidity and mortality, particularly among people with advanced HIV disease and others with impaired cell-mediated immunity. Primarily caused by *Cryptococcus neoformans*, it is a leading cause of life-threatening fungal meningitis in immunocompromised individuals, responsible for over 120,000 deaths annually and imposing an economic burden of >$535 million in the United States alone (1–6). Recently, the WHO recognized *C. neoformans* as a high-priority fungal pathogen based on its global burden, relapse rates, and limited therapeutic options. Because antifungal therapy alone rarely resolves cryptococcal disease, outcomes largely depend on the quality of the host immune response, making immunology-focused investigations crucial for developing new therapies.

Neutrophils are the most abundant leukocyte in human circulation and the earliest responders to most microbial challenges, equipped with reactive oxygen species, granule proteases, antimicrobial peptides, and extracellular traps. For most invasive fungal infections, they are indispensable. Neutropenia is among the strongest clinical risk factors for invasive candidiasis and aspergillosis, and inherited defects in neutrophil number or effector function predispose to severe, often fatal mold and yeast disease (7, 8). However, their role in cryptococcosis remains paradoxical and poorly defined. Neutropenia is not a recognized risk factor for cryptococcal disease, and among patients with cryptococcal meningitis, elevated neutrophil-to-lymphocyte ratios predict unfavorable outcomes (9, 10). These observations have supported the prevailing view that neutrophils are dispensable, or even deleterious, during cryptococcal infection. However, neutrophils appear to have context-dependent roles during cryptococcosis, with neutrophils shown to worsen or improve survival depending on the model, fungal strain, host background, and timing of depletion (11–14). In a vaccine model, neutrophil depletion can eliminate protection, further supporting context-dependent protective functions. These contradictions likely reflect neutrophil functional heterogeneity, as distinct neutrophil subsets with divergent transcriptional programs are generated during cryptococcal lung infection (15). Neutrophils are also among the earliest leukocytes recruited to the lung after cryptococcal infection, peaking before the accumulation of monocytes and other inflammatory cells (16), positioning them to shape subsequent immune responses.

Beyond direct antifungal activity, neutrophils are increasingly recognized as immunomodulators that shape the function of other immune cells (17–21). For example, in *Aspergillus fumigatus* infection, neutrophils “license” monocytes for fungicidal activity, and without them, monocytes fail to mature or function effectively (22). We have shown in a related cryptococcal model that neutrophils are not intrinsically pathogenic but instead mediate lethal immunopathology through reactive oxygen species-dependent amplification of inflammation (23). Together, these findings suggest that neutrophils in cryptococcosis extend beyond direct antifungal activity to regulate broader host immune responses.

Here, we investigated the role of neutrophils during pulmonary infection with *C. neoformans* using complementary Ly6G-specific depletion approaches. Contrary to prior reports, we found that neutrophil depletion markedly accelerates mortality without changing fungal burden. This phenotype is independent of mouse genetic background, depletion strategy, inoculum, and fungal strain, and is reproduced with phenotypically divergent clinical isolates. Cytokine profiling and immunophenotyping reveal that neutrophil loss provokes an early chemokine and IL-17A response, followed by massive accumulation of inflammatory monocytes and a reciprocal collapse of type 2 cytokines, eosinophils, and alveolar macrophages. Most notably, the increased mortality associated with neutrophil depletion was completely absent in CCR2-deficient mice. These data identify neutrophils as critical immunoregulatory cells in cryptococcosis that restrain pathogenic, CCR2-dependent monocyte-driven inflammation, and they redefine the contribution of neutrophils to host defense against *C*. *neoformans*.

## RESULTS

### Neutrophil-depleted mice succumb to infection more rapidly

It has been previously reported that neutrophil depletion leads to increased survival of *C. neoformans-infected* mice (11). To define the role of neutrophils in the *C. neoformans* immune response, we first depleted neutrophils in male and female C57BL/6 mice by injecting 200 µg of α-Ly6G (1A8 (24, 25)) or an isotype antibody, followed 24h later by inoculation with 10^5^ KN99 via orotracheal aspiration. Antibodies were administered every 48 hours, and mice were monitored for survival, with humane endpoints defined as ≥20% weight loss or morbidity.

Contrary to prior reports, neutrophil-depleted mice succumbed more rapidly than controls (Fig. 1A), with a sharp decline in body weight, resulting in early morbidity and euthanasia (Fig. 1B). These findings were unexpected and suggest that, under our experimental conditions, neutrophils play a protective rather than detrimental role during *Cn* infection. We next tested whether the accelerated mortality of neutrophil-depleted mice was due to differences in fungal burden. Male and female C57BL/6 mice were depleted and infected as described above. We harvested lungs, brains, and spleens at 7d post-infection, when neutrophil-depleted mice began to succumb (Fig. 1A), and plated the homogenates on YPD plates for CFU quantification. We found no significant differences in fungal burden in the lungs, brain, or spleen (Fig. 1C) at any time point. These findings suggest that accelerated mortality in neutrophil-depleted mice is due to dysregulated host immunity rather than unchecked fungal growth.

**Figure 1.**
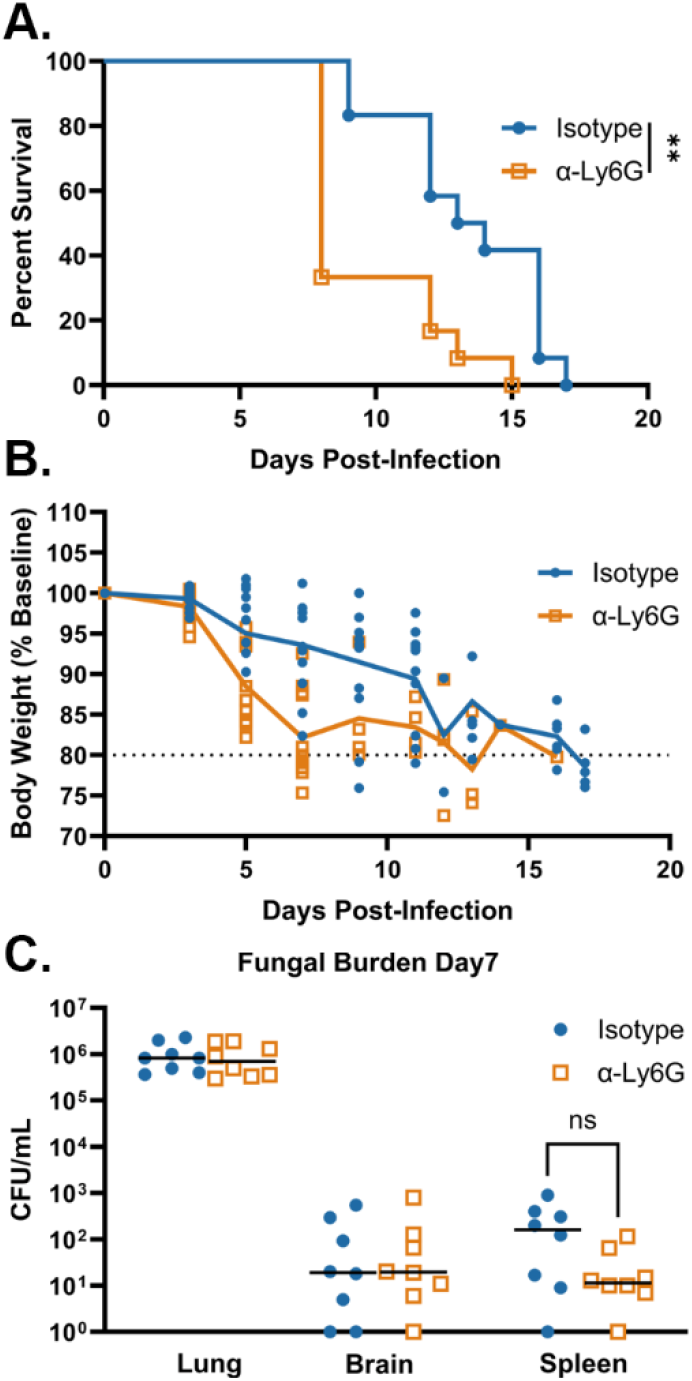
Neutrophil depletion accelerates mortality without altering fungal burden. Male and female C57BL/6 mice received 200 µg of anti-Ly6G (clone 1A8) or rat IgG2a isotype control antibody by intraperitoneal injection beginning 24 h before infection and every 48 h thereafter and were inoculated by orotracheal aspiration with 10^5^ CFU of *C. neoformans* KN99. (A) Survival. Mice that lost 20% of the body weight recorded at the time of inoculation or displayed signs of morbidity were considered ill and euthanized. (B) Body weight, expressed as a percentage of the weight recorded at inoculation. The dotted line indicates the 20% weight-loss endpoint. (C) Fungal burden in the lung, brain, and spleen at 7 days post-infection. Each symbol represents an individual mouse. Data are cumulative of three independent experiments with a total of 12 mice per group (A and B) and of two independent experiments with a total of 8 per group (C). **, *P* < 0.01; ns, not significant.

### Mortality is not due to antibody, mouse background, inoculum, or strain of C. neoformans

We observed a phenotype different from what was previously reported. Several factors may explain this discrepancy, including differences in mouse backgrounds, antibody specificity (the original antibody also depleted monocytes), dosing regimen, and fungal strain, as we used the highly virulent KN99 (11). To determine whether our findings were influenced by the depletion antibody or mouse genetics, we employed a recently developed recombinant murinized α-Ly6G (1A8-CP129) antibody, which achieves robust neutrophil depletion across multiple mouse strains and overcomes limitations of conventional rat-derived antibodies (26). To confirm depletion efficiency, C57BL/6 mice were injected with 100 µg of 1A8-CP129 or isotype control, and neutrophils were quantified in blood, lungs, and spleens at 48h. 1A8-CP129 efficiently depleted neutrophils in all three compartments, reducing neutrophils from 7.03 ± 0.10% to 0.08 ± 0.01% of CD45^+^ cells in the lung, from 6.58 ± 0.48% to 0.43 ± 0.08% in blood, and from 1.83 ± 0.27% to 0.01 ± 0.00% in spleen (n = 3 per group; p < 0.01 for each compartment), corresponding to a 98.8–99.9% reduction in absolute neutrophil number at every site (Fig. 2A). Next, C57BL/6, BALB/c, and CBA/J mice were treated with 1A8-CP129 or isotype control and infected as described above. Across all strains tested, neutrophil-depleted mice succumbed to infection more rapidly than controls (Fig. 2B-D), demonstrating that accelerated mortality is independent of mouse background and depletion strategy. Accelerated mortality was also independent of inoculum. Neutrophil-depleted mice succumbed more rapidly than isotype-treated controls after challenge with either 10⁴ or 10⁵ CFU of KN99 (Fig. S1).

**Figure 2.**
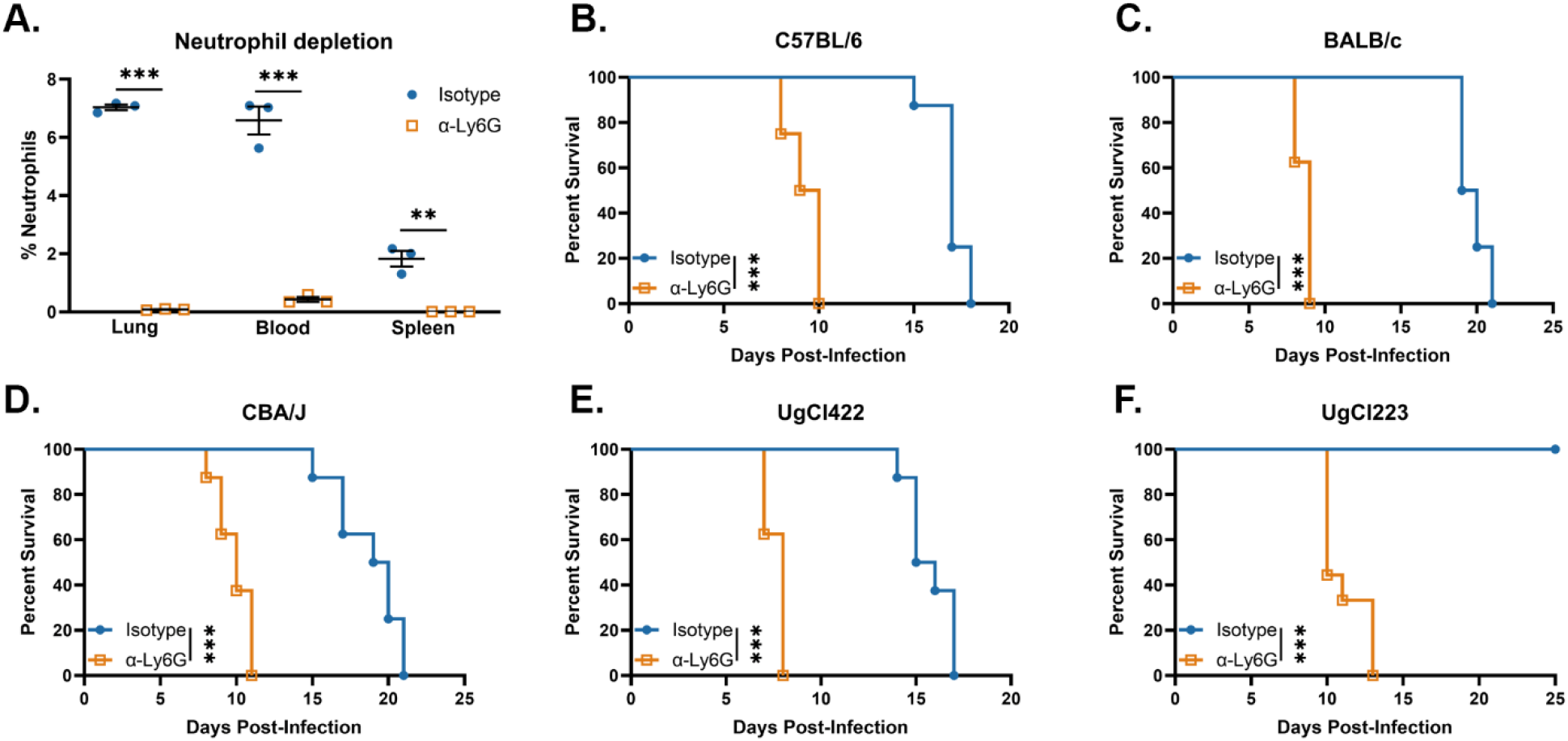
Accelerated mortality in neutrophil-depleted mice is independent of depletion antibody, mouse genetic background, and fungal strain. (A) Depletion efficiency. C57BL/6 mice received a single intraperitoneal injection of 100 µg of recombinant murinized anti-Ly6G (1A8-CP129) or isotype control antibody, and neutrophils were quantified in the lung, blood, and spleen by flow cytometry 48 h later. Values are expressed as a percentage of CD45^+^ cells. Bars indicate means ± standard errors of the means (SEM), and each symbol represents an individual mouse (*n* = 3 per group). (B to D) C57BL/6 (B), BALB/c (C), and CBA/J (D) mice were treated with 1A8-CP129 or isotype control and inoculated with 10^5^ CFU of KN99. (E and F) C57BL/6 mice were treated as described for panels B to D and inoculated with 10^5^ CFU of the hypervirulent clinical isolate UgCl422 (E) or the hypovirulent clinical isolate UgCl223 (F). Survival data are cumulative of two independent experiments with a total of 8 mice per group. **, *P* < 0.01; ***, *P* < 0.001.

We used the highly virulent KN99 for our assays, however, the original neutrophil depletion paper used a less virulent strain (11). To determine whether this phenotype was restricted to KN99, C57BL/6 mice were depleted of neutrophils using 1A8-CP129 or isotype control and infected with 10⁵ cells of either the hypovirulent UgCl223 or the hypervirulent UgCl422 clinical isolate(27, 28). Consistent with our findings using KN99, neutrophil-depleted mice infected with either clinical isolate succumbed significantly more rapidly than isotype-treated controls (Figs. 2E and 2F), demonstrating that neutrophil-dependent protection is conserved across phenotypically diverse clinical isolates of *C. neoformans* and is not attributable to differences in intrinsic fungal virulence. We used KN99 and the 1A8-CP129 antibody for all following experiments.

### Increased proinflammatory cytokines underlie accelerated mortality in neutrophil-depleted mice

Our data indicate that the increased mortality of neutrophil-depleted mice is not due to differences in fungal burden (Fig. 1C), suggesting that death is host-mediated. To begin characterizing the host response, we analyzed cytokines and chemokines from the lungs of depleted and control mice. Male and female C57BL/6 mice were depleted and infected as described above, and lungs were harvested on day 7. Tissue homogenates were analyzed for cytokine/chemokine responses using a 26-plex cytokine and chemokine panel. Thirteen of 26 analytes differed significantly at day 7, whereas none of the 26 differed between isotype- and 1A8-CP129-treated mice in the absence of infection (Fig. 3A), indicating that neutrophil depletion alone does not perturb the pulmonary cytokine milieu. On day 7, neutrophil-depleted mice exhibited a marked increase in proinflammatory cytokines, with MIP-2α/CXCL2 increased 6.8-fold, TNF-α 5.7-fold, and IL-6 3.6-fold (Figs. 3B-D). This was accompanied by a significant decrease in the type 2 cytokines IL-4, IL-5, and IL-13 (Figs. 3E-G), which are generally associated with poor outcomes in cryptococcal infection (29, 30). These findings indicate that neutrophil-depleted mice mount a heightened proinflammatory immune response that is amplified by suppression of Th2 cytokines.

**Figure 3.**
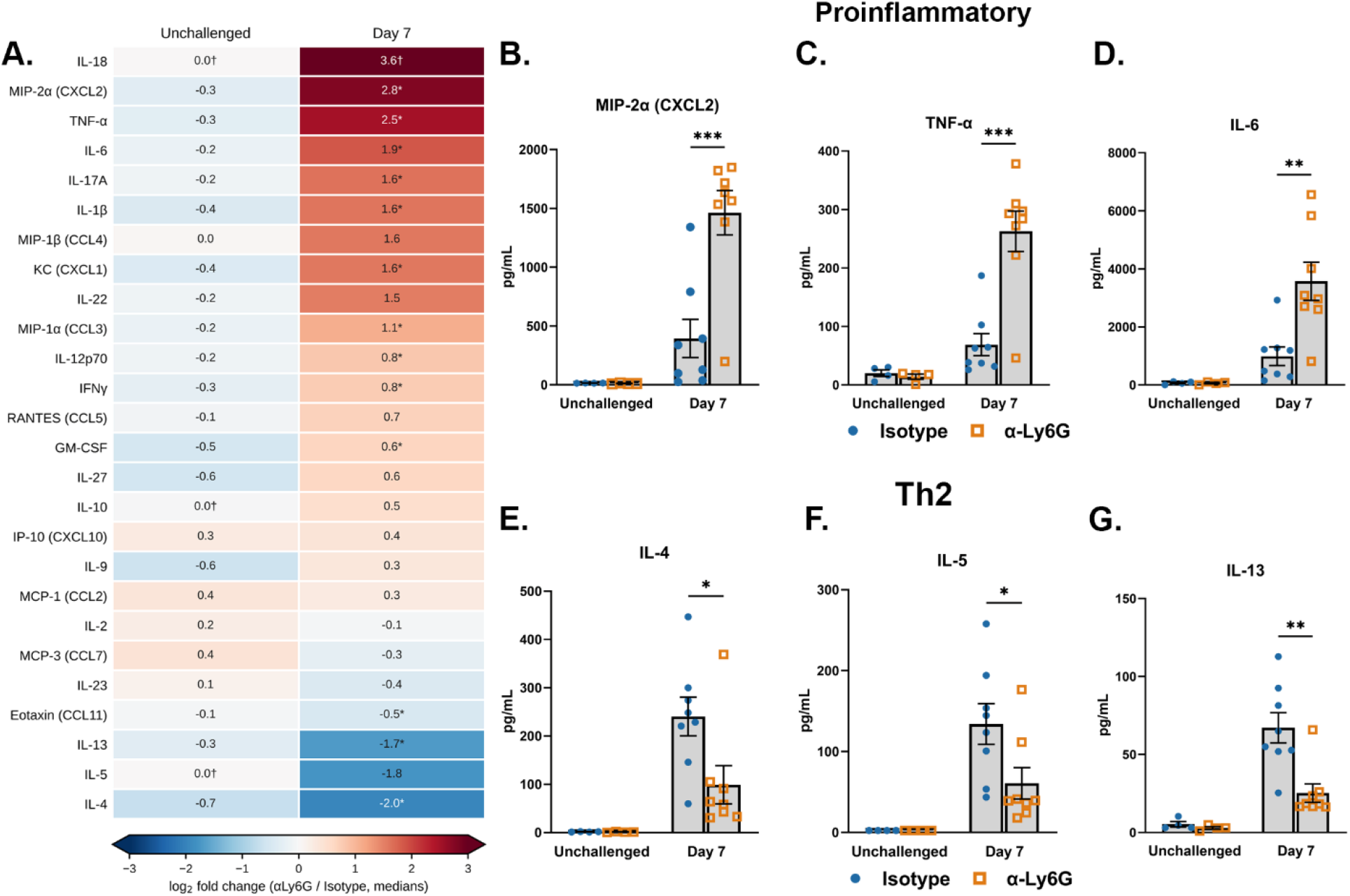
Neutrophil depletion amplifies proinflammatory cytokines and suppresses type 2 cytokines in the lung at day 7. Male and female C57BL/6 mice were depleted of neutrophils with 1A8-CP129 or given isotype control antibody and inoculated with 10^5^ CFU of KN99. Lungs were harvested from unchallenged animals and at 7 days post-infection, and homogenates were assayed with a 26-plex cytokine and chemokine panel. (A) Heat map of all 26 analytes, ordered by the magnitude of the day 7 response. Values are log_2_ fold changes in group medians (anti-Ly6G/isotype); red indicates higher and blue lower concentrations in neutrophil-depleted mice, as shown in the color scale. (B to D) Proinflammatory analytes MIP-2α/CXCL2 (B), TNF-α (C), and IL-6 (D). (E to G) Type 2 analytes IL-4 (E), IL-5 (F), and IL-13 (G). Bars indicate means ± SEM, and each symbol represents an individual mouse. Data are cumulative from one experiment for unchallenged mice, for a total of 4 per group, and two independent experiments for day 7, with a total of 8 mice per group. *, *q* < 0.05; **, *q* < 0.01; ***, *q* < 0.001. †, 25% or more of the values in at least one group fell at the assay lower limit of detection. Color-capped at ± 3; printed values are not.

### Flow cytometry of the pulmonary myeloid compartment on day 7 revealed a dominant inflammatory monocyte infiltrate

To further characterize the host response, we analyzed the cellular profiles from the lungs of depleted and control mice. We depleted and infected C57BL/6 mice as described above, harvested lungs on day 7, and isolated pulmonary cells for flow cytometry. We observed that total CD45^+^ numbers were unchanged (Fig. 4A), and neutrophil depletion remained efficient at this time point, with a 93% reduction in pulmonary neutrophils (Fig. 4B). We observed a significant increase in monocytes and inflammatory macrophages in the depleted mice compared to controls, with Ly6C^high^ monocytes increased 11.5-fold and Ly6C^int^ monocytes 9.4-fold, with inflammatory macrophages increased 3.4-fold (Figs. 4B, C, and F). Two populations were significantly reduced in depleted mice compared with controls. Alveolar macrophages fell by 89% (Fig. 4H) and eosinophils by 72% (Fig. 4I), with the latter accompanied by reduced eotaxin/CCL11 (Fig. 3A). In contrast, no lymphocyte population differed between groups, including CD4^+^ and CD8^+^ T cells, γδ T cells, NK cells, NKT cells and B cells (Fig. S2), indicating that the day 7 response to neutrophil depletion is confined to the myeloid compartment. Additionally, we did not observe any difference in CD11b or monocyte Dendritic cell populations (Fig. S2). Taken together, these findings suggest that in the absence of neutrophils, excessive recruitment and activation of monocytes amplify a hyperinflammatory but non-protective immune response, explaining the accelerated mortality we observed.

**Figure 4.**
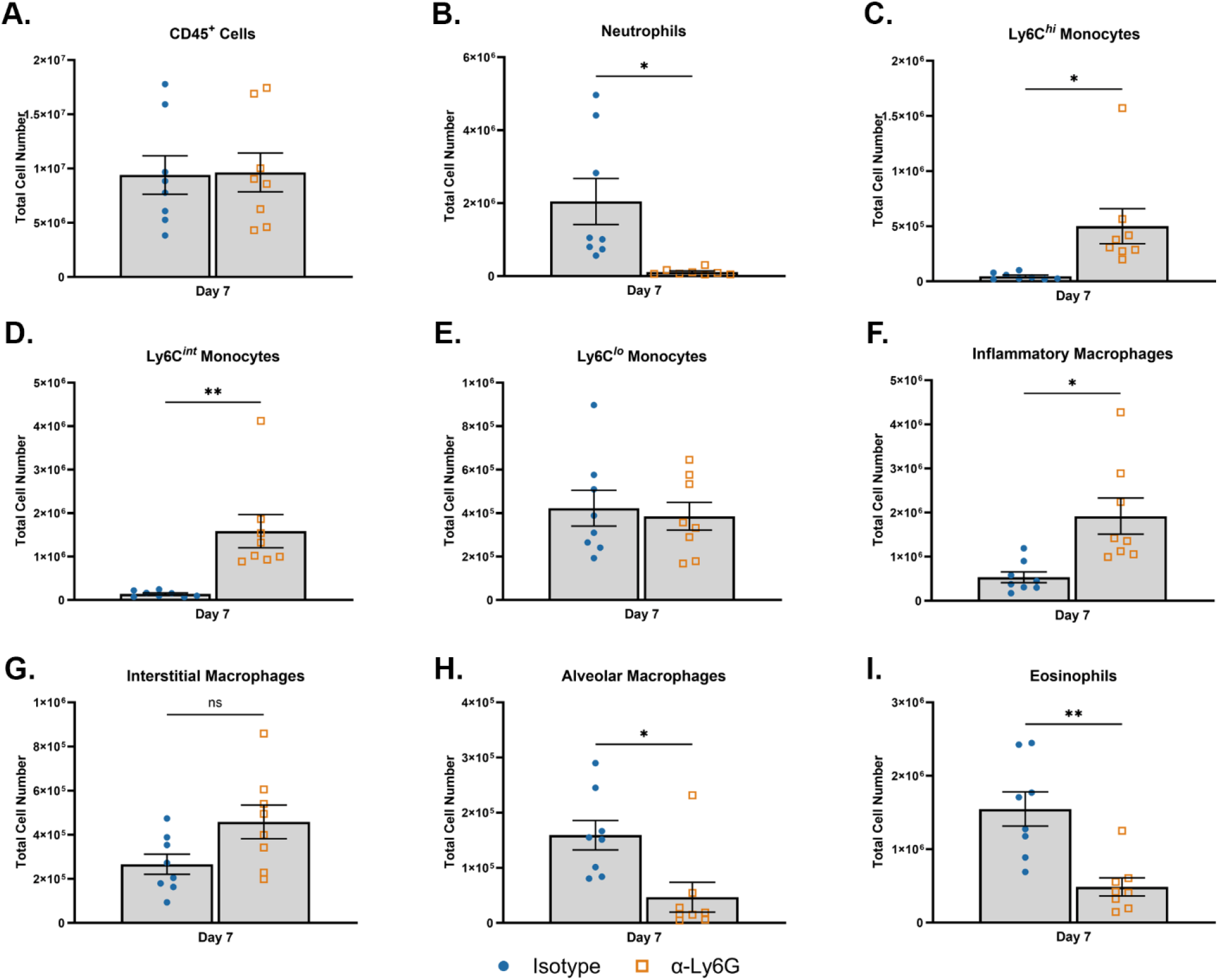
Neutrophil depletion produces a dominant inflammatory monocyte infiltrate in the lung at day 7. Male and female C57BL/6 mice were depleted of neutrophils with 1A8-CP129 or given isotype control antibody and inoculated with 10^5^ CFU of KN99. Lungs were harvested at 7 days post-infection, and pulmonary leukocytes were isolated and subjected to flow cytometric analysis (see Table S1 for antibodies and Fig. S3 for the gating strategy). Total numbers are shown for CD45^+^ leukocytes (A), neutrophils (B), Ly6C^hi^ monocytes (C), Ly6C^int^ monocytes (D), Ly6C^lo^ monocytes (E), inflammatory macrophages (F), interstitial macrophages (G), alveolar macrophages (H), and eosinophils (I). Bars indicate means ± SEM, and each symbol represents an individual mouse. Data are cumulative of two independent experiments with a total of 8 mice per group. *, *P* < 0.05; **, *P* < 0.01; ns, not significant.

### Neutrophil depletion provokes an early chemokine and IL-17A response

Neutrophils are among the earliest cell types recruited to the lungs during *C. neoformans* infection, peaking approximately 18 h before declining as other immune cells are recruited (16). Because this peak precedes day 7 by nearly a week, we examined early time points in addition to day 7, reasoning that the interval in which neutrophils are present is the one most likely to reveal how their loss shapes the subsequent response. Male and female C57BL/6 mice were depleted and infected as described above, and lungs were harvested from unchallenged animals and at 24h and 48h post-infection. Tissue homogenates were analyzed for cytokine/chemokine responses using a 26-plex cytokine and chemokine panel. Ten of 26 analytes differed significantly between groups (Fig. 5A). The early response was dominated by neutrophil- and monocyte-directed chemokines, along with IL-17A, rather than by classical proinflammatory cytokines. KC/CXCL1 was the most strongly induced analyte, increasing 121-fold and remaining elevated at 48 h (Fig. 5B). IL-17A increased approximately 14-fold at 24 h and 15-fold at 48 h (Fig. 5C), and IL-6 rose 5.9-fold at 24 h and 9.7-fold at 48 h (Fig. 5D). Additionally, the monocyte-recruiting chemokines MIP-1α/CCL3, MIP-1β/CCL4, MCP-1/CCL2, MCP-3/CCL7 and IP-10/CXCL10 were all significantly elevated in the depleted mice (Fig. 5A, E). The co-induction of KC/CXCL1 and IL-17A is consistent with the IL-17-dependent granulopoietic feedback that is engaged when circulating neutrophils are cleared (31). The concurrent elevation of CCL2, CCL3, CCL4 and CCL7 provides a chemotactic basis for the monocyte accumulation observed at day 7 (Fig. 4).

**Figure 5.**
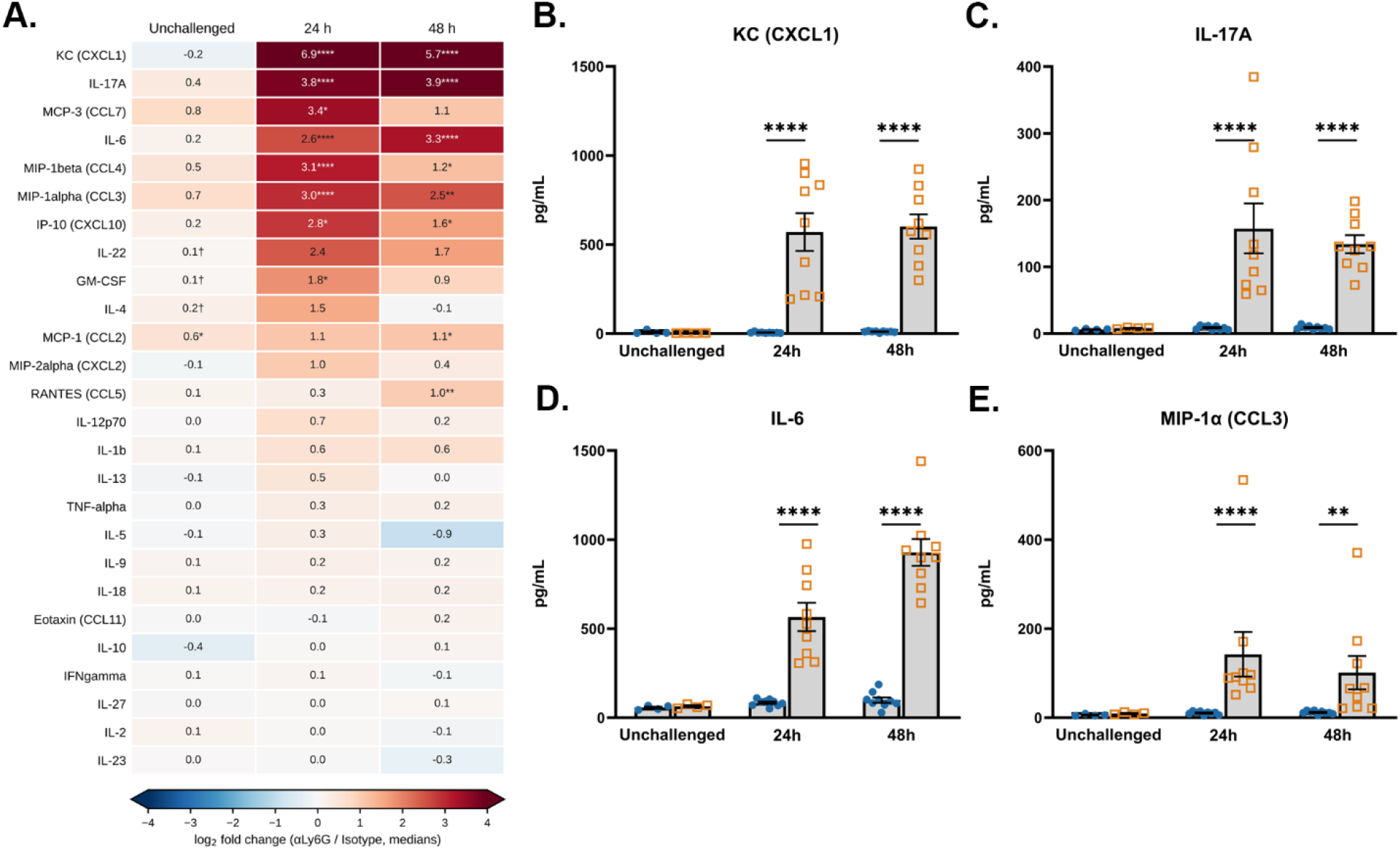
Neutrophil depletion provokes an early pulmonary chemokine and IL-17A response. Male and female C57BL/6 mice were depleted of neutrophils with 1A8-CP129 or given isotype control antibody and inoculated with 10^5^ CFU of KN99. Lungs were harvested from unchallenged animals and at 24 and 48 h postinfection, and homogenates were assayed with a 26-plex cytokine and chemokine panel. (A) Heat map of all 26 analytes, ordered by the magnitude of the 24-h response. KC/CXCL1 (B), IL-17A (C), IL-6 (D), and MIP-1α/CCL3 (E). Bars indicate means ± SEM, and each symbol represents an individual mouse. Data are cumulative from one experiment for unchallenged mice, for a total of 4 per group, and two independent experiments with a total of 9 mice per group per time point. *, *q* < 0.05; **, *q* < 0.01; ****, *q* < 0.0001. †, 25% or more of the values in at least one group fell at the assay lower limit of detection. Color-capped at ± 4; printed values are not.

Changes in cytokine and chemokine levels could reflect differences in fungal burden rather than altered host regulation. Although burden was indistinguishable between groups at day 7 (Fig. 1C), this does not exclude an earlier divergence that could account for the responses observed at 24h and 48h. Male and female C57BL/6 mice were therefore depleted and infected as described above, lungs were harvested at 24h and 48h post-infection, and homogenates were plated on YPD agar for CFU quantification. We found no significant differences in fungal burden in the lungs at any time point (Fig. 6A). Together, these findings demonstrate that the heightened inflammatory response and accelerated mortality associated with neutrophil depletion occur despite comparable pulmonary fungal burdens at 24h and 48h (Fig. 6A), and day 7 post-infection (Fig. 1C), supporting a role for dysregulated immunity as a major driver of disease severity.

**Figure 6.**
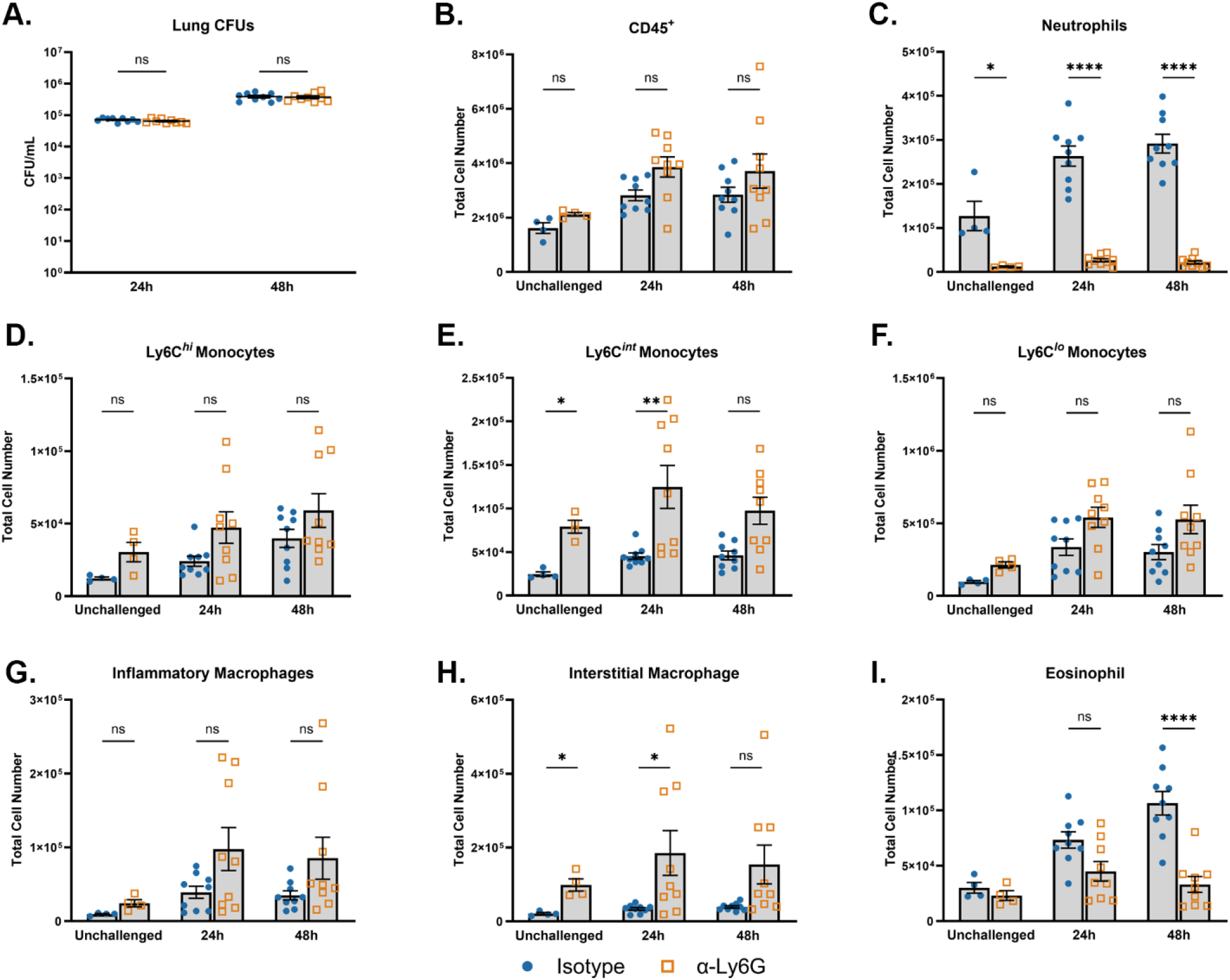
Neutrophil depletion remodels the pulmonary myeloid compartment within 24h without altering fungal burden. Male and female C57BL/6 mice were depleted of neutrophils with 1A8-CP129 or given isotype control antibody and inoculated with 10^5^ CFU of KN99. (A) Pulmonary fungal burden at 24 and 48h post-infection. Each symbol represents an individual mouse. (B to I) Pulmonary leukocytes were isolated from unchallenged animals and at 24 and 48 h post-infection and subjected to flow cytometric analysis. Total numbers are shown for CD45^+^ leukocytes (B), neutrophils (C), Ly6C^hi^ monocytes (D), Ly6C^int^ monocytes (E), Ly6C^lo^ monocytes (F), inflammatory macrophages (G), interstitial macrophages (H), and eosinophils (I). Bars indicate means ± SEM, and each symbol represents an individual mouse. Data are cumulative of two independent experiments with a total of 9 mice per group at 24 and 48h and 4 mice per group for unchallenged controls. *, *P* < 0.05; **, *P* < 0.01; ****, *P* < 0.0001; ns, not significant.

### Neutrophil depletion remodels the pulmonary myeloid compartment within 24 h of infection

To determine whether the increased production of neutrophil- and monocyte-associated chemokines corresponded with changes in immune cell recruitment, we characterized pulmonary leukocyte populations by flow cytometry. Male and female C57BL/6 mice were depleted of neutrophils and infected as described above. Lungs were harvested from unchallenged mice and at 24h and 48h post-infection for flow cytometric analysis. Total pulmonary CD45⁺ cell numbers were comparable between groups at all time points (Fig. 6B), despite effective neutrophil depletion, which reduced pulmonary neutrophil numbers by 91% at 24h and 93% at 48h post-infection (Fig. 6C). Among the myeloid populations examined, Ly6Ci^nt^ monocytes and interstitial macrophages exhibited the most consistent changes, with both populations significantly increased in neutrophil-depleted mice under unchallenged conditions and at 24h post-infection (Figs. 6E and 6H). In contrast to the significant expansion observed at day 7, Ly6C^high^ monocytes did not differ significantly between groups at these early time points, although an upward trend was evident in neutrophil-depleted mice (Fig. 6D). Conversely, pulmonary eosinophil numbers were reduced by 72% at 48h post-infection (Fig. 6I), consistent with the reductions observed at day 7 (Fig. 4I). Collectively, these findings demonstrate that neutrophil depletion rapidly reshapes the pulmonary myeloid compartment, characterized by early accumulation of Ly6C^int^ monocytes and interstitial macrophages alongside a marked reduction in eosinophils, preceding the more pronounced inflammatory changes observed later during infection.

### CCR2 deficiency abrogates the accelerated mortality associated with neutrophil depletion

To identify the immune cell populations responsible for the accelerated mortality observed following neutrophil depletion, we first investigated the contribution of adaptive immunity.

Given the strong association between cryptococcosis and impaired CD4⁺ T cell immunity, we examined whether the increased mortality in neutrophil-depleted mice was dependent on B or T cells. Neutrophil-depleted Rag1^⁻/⁻^ mice, which lack mature B and T lymphocytes, exhibited accelerated mortality comparable to that observed in neutrophil-depleted wild-type (WT) mice (Fig. 7A). This finding was consistent with our day 7 flow cytometric analysis, which revealed no significant differences in pulmonary adaptive immune cell populations between neutrophil-depleted and control mice (Fig. S2). Together, these results indicate that adaptive immunity is not required for the increased susceptibility associated with neutrophil depletion, suggesting that dysregulation of innate immune responses is primarily responsible for the observed phenotype.

**Figure 7.**
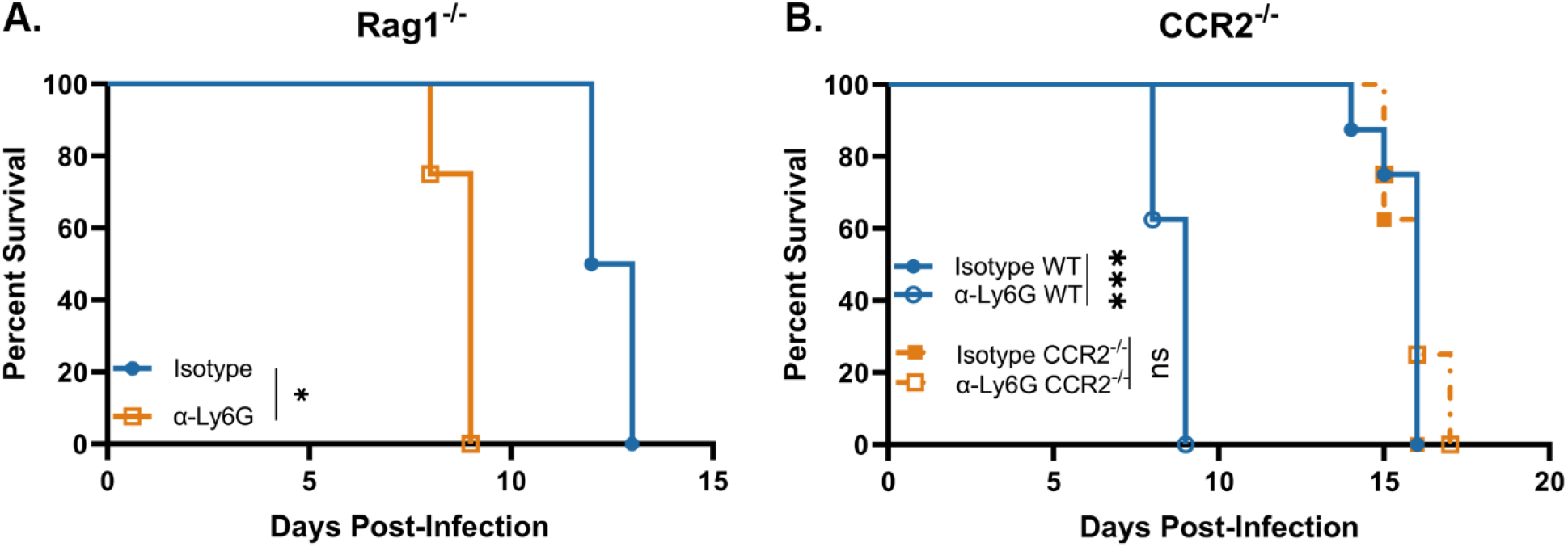
Accelerated mortality in neutrophil-depleted mice requires CCR2 but not adaptive immunity. (A) *Rag1*^-/-^ mice, which lack mature B and T lymphocytes, were treated with 1A8-CP129 or isotype control antibody and inoculated with 10^5^ CFU of KN99. (B) Wild-type (WT) C57BL/6 and *Ccr2*^-/-^ mice were treated and infected in parallel. The four groups shown are isotype-treated WT, anti-Ly6G-treated WT, isotype-treated *Ccr2*^-/-^, and anti-Ly6G-treated *Ccr2*^-/-^ mice. Mice that lost 20% of the body weight recorded at the time of inoculation or displayed signs of morbidity were considered ill and euthanized. Data are cumulative of two independent experiments with a total of 8 mice per group. *, *P* < 0.05; ***, *P* < 0.0001; ns, not significant.

Given the pronounced accumulation of inflammatory monocytes in neutrophil-depleted mice, we next investigated whether altered monocyte trafficking contributed to accelerated mortality. Monocytes play complex roles during cryptococcosis, contributing to antifungal defense while also promoting detrimental inflammation under certain conditions(32). In particular, excessive CCR2⁺ Ly6C^high^ monocyte recruitment has been shown to worsen disease outcomes by fueling tissue-damaging inflammation and supporting fungal persistence (33). Furthermore, monocyte dysfunction in patients with cryptococcal meningitis has been strongly correlated with mortality (34). To determine whether the accelerated mortality of neutrophil-depleted mice was driven by excessive monocyte recruitment, we used CCR2-deficient mice (*Ccr2^-/^*^-^) (35), which cannot mobilize Ly6C^high^ inflammatory monocytes from the bone marrow and display ∼75% fewer circulating monocytes under homeostatic conditions (33, 36). Male and female *Ccr2^-/^*^-^ and wild-type (WT) controls were depleted of neutrophils and infected as described above.

Strikingly, whereas neutrophil depletion significantly accelerated mortality in WT mice, this effect was completely absent in *Ccr2^-/^*^-^ mice, with neutrophil-depleted *Ccr2^-/^*^-^ animals exhibiting survival comparable to that of both isotype-treated control groups (Fig. 7B). These findings establish that CCR2-dependent immune responses are critical for the increased mortality associated with neutrophil depletion and strongly implicate inflammatory monocytes as key mediators of disease severity. Collectively, our results identify a previously unrecognized protective function of neutrophils during cryptococcal infection, whereby they restrain CCR2-dependent, monocyte-associated immunopathology.

## DISCUSSION

The longstanding view that neutrophils are dispensable during cryptococcosis has been shaped largely by the fact that neutropenia is not a recognized clinical risk factor for cryptococcal disease, and an early murine study demonstrated that neutrophil depletion reduced susceptibility to pulmonary infection (11). Our findings challenge this interpretation and reveal a previously underappreciated protective role for neutrophils during pulmonary cryptococcosis. Across three mouse genetic backgrounds, two neutrophil depletion strategies, two infectious inocula, and three *C. neoformans* strains with differing virulence, neutrophil depletion consistently accelerated mortality. Importantly, this increased mortality occurred without corresponding increases in pulmonary fungal burden at 24h, 48h, or 7 days post-infection, or in the brain and spleen at day 7. These findings suggest that the accelerated mortality results from dysregulated host inflammation rather than impaired fungal control. Immunophenotyping localized the dysregulated response almost entirely to the myeloid compartment, identified a dominant inflammatory monocyte infiltrate, and the complete rescue in *Ccr2*^-/-^ mice establishes CCR2-dependent monocyte recruitment as the proximate cause of death. In this setting, neutrophils function less as antifungal effectors than as regulators that constrain a lethal monocyte response.

Several methodological differences likely reconcile our results with the original depletion study. That work used the anti-Gr-1 antibody RB6-8C5, which recognizes both Ly6G and Ly6C and therefore depletes monocytes, dendritic cells, and lymphocyte subsets in addition to neutrophils, whereas the Ly6G-specific clone 1A8 depletes neutrophils while sparing Gr-1^+^ monocytes (37). Because our data identify CCR2^+^ monocytes as the effectors of mortality, an antibody that removes neutrophils and monocytes concurrently would eliminate precisely the population responsible for the pathology that neutrophil-selective depletion unmasks. This could explain why broad Gr-1 depletion improved survival while Ly6G-specific depletion worsens it, and it is consistent with the observation that RB6-8C5 pretreatment amplifies systemic TNF-α responses far more than 1A8 (37). Fungal virulence also differs, as the original study used a less virulent isolate than the hypervirulent KN99 strain used here (11). To exclude antibody- and host-specific artifacts, we employed the recombinant murinized 1A8-CP129 antibody, which achieved 98.8-99.9% reductions in absolute neutrophil number in lung, blood, and spleen and overcomes the anti-rat immune responses that limit conventional rat-derived clones (26). The phenotype persisted in C57BL/6, BALB/c, and CBA/J mice and in animals infected with the hypovirulent UgCl223 and hypervirulent UgCl422 clinical isolates (27, 28) (Fig. 2), arguing that neutrophil-dependent immune regulation is a general feature of pulmonary cryptococcosis rather than a property of a single strain or genetic background.

Prior conclusions that neutrophils are unimportant in cryptococcosis have largely been drawn from measurements of fungal clearance. We found that neutrophils are not required to control cryptococcal growth, consistent with reports that antibody depletion of neutrophils does not alter pulmonary fungal clearance (38). A limited role in fungal clearance does not diminish the importance of neutrophils for host survival. Rather, our findings suggest that neutrophils protect against cryptococcosis primarily by regulating host inflammatory responses, a function reflected in survival outcomes despite unchanged fungal burden. This interpretation parallels our previous observation that neutrophils recruited to the lung in the absence of NADPH oxidase 2 remain abundant yet harmless, demonstrating that neutrophil number and neutrophil consequence can be fully uncoupled (23). Single-cell studies identifying transcriptionally distinct neutrophil populations during cryptococcal infection offer one possible explanation for such functional diversity (15).

The kinetics of the response suggest how neutrophil loss is translated into monocyte-driven pathology. Because pulmonary neutrophils peak near 18h post-infection (16), we examined the interval in which they are normally present. Within 24h of infection, the lungs of neutrophil-depleted mice were dominated not by classical proinflammatory cytokines but by chemokines and IL-17A. KC/CXCL1 was the most strongly induced analyte, rising 121-fold, and IL-17A increased approximately 14-fold (Fig. 5). The striking increase in IL-17A merits particular attention because neutrophil abundance is subject to an IL-23–IL-17-mediated homeostatic feedback circuit (31). In a protective cryptococcal immunization model, it was similarly found that neutrophil depletion increased IL-17A, IL-6, and CXCL1, with γδ T cells contributing to the IL-17A response (39). Concurrently, the monocyte-recruiting chemokines CCL2, CCL3, CCL4, CCL7, and CXCL10 were all significantly elevated, providing a direct chemotactic basis for the myeloid remodeling that followed. Ly6C^int^ monocytes and interstitial macrophages increased as early as 24h, and by day 7 Ly6C^high^ and Ly6C^int^ monocytes had expanded 11.5-fold and 9.4-fold with a 3.4-fold increase in inflammatory macrophages. Notably, neutrophil depletion alone perturbed none of the 26 analytes in uninfected animals, indicating that this cascade requires the combination of infection and neutrophil absence rather than depletion itself.

That CCR2 deficiency completely abrogated the excess mortality places inflammatory monocytes at the end of this pathway. *Ccr2*^-/-^ mice cannot mobilize Ly6C^high^ monocytes from the bone marrow and circulate approximately 75% fewer monocytes at homeostasis (35, 36), and in these animals neutrophil depletion conferred no survival disadvantage whatsoever. This result is consistent with a growing body of work establishing that monocyte responses in cryptococcosis are double-edged. CCR2 signaling governs the recruitment of monocytes and conventional DCs and the formation of bronchovascular mononuclear infiltrates during cryptococal infection (40), and excessive CCR2^+^ Ly6C^high^ monocyte recruitment worsens outcome by fueling tissue damage and supporting fungal persistence (32, 33). Monocytes can promote antifungal defense through phagocytosis, cytokine production, and differentiation into type 1 macrophages and DCs, yet *C. neoformans* can co-opt them toward type 2 and alternatively activated programs that favor persistence and dissemination (41–45). Our data add an upstream layer to this picture by identifying neutrophils as the constraint that keeps CCR2^+^ monocyte responses in check.

How neutrophils restrain monocytes in this setting remains to be defined. Neutrophils can interact with monocytes, macrophages, and dendritic cells through both contact-dependent and soluble mechanisms, as exemplified by Mac-1-dependent suppression of lymphocyte responses, extracellular trap-mediated dendritic cell activation, granule proteins, and extracellular vesicles (21, 46, 47), and the licensing of monocytes by neutrophils during aspergillosis demonstrates that such crosstalk can determine the functional maturation of the monocyte compartment (22). Our data are most consistent with the loss of an active brake rather than the loss of a maturation signal, since monocytes in neutrophil-depleted mice were not deficient but excessive and hyperinflammatory.

Future studies will seek to determine whether neutrophils limit inflammatory monocyte recruitment, activation, or both, and whether these regulatory effects are mediated through direct cell–cell interactions or soluble factors. Co-culture assays, adoptive transfer experiments, and transcriptional profiling of monocytes isolated from neutrophil-depleted and control mice will help define the mechanisms underlying this regulation. Because germline *Ccr2*^-/-^ mice exhibit altered monocyte homeostasis and impaired mobilization from the bone marrow, the protection observed in these animals cannot be attributed solely to reduced monocyte recruitment (48). Pharmacologic CCR2 blockade and adoptive monocyte transfer studies will therefore be important for establishing the necessity and sufficiency of inflammatory monocytes in mediating disease pathology and determining whether these cells initiate or amplify the dysregulated inflammatory response.

Our data establish that neutrophils are not dispensable in cryptococcosis. They reframe neutrophils not as antifungal effectors whose absence is tolerated, but as upstream regulators of monocyte-driven immunopathology whose loss converts a controlled infection into a lethal inflammatory disease, and they identify the CCR2 axis as the effector arm of that pathology and a rational target for host-directed therapy.

## MATERIAL AND METHODS

### Ethics Statement

All animal experiments were conducted in compliance with the Public Health Service Policy on Humane Care and Use of Laboratory Animals and adhered to the guidelines in the Guide for the Care and Use of Laboratory Animals. The study protocols were reviewed and approved by the Institutional Animal Care and Use Committee (IACUC) at the University of Tennessee Health Science Center (UTHSC) under protocol number 24-0556. All mice were housed in AAALAC-accredited facilities located at UTHSC under conditions of controlled temperature, humidity, and a 12-hour light/dark cycle. Mice were provided standard rodent chow and water *ad libitum* and were monitored daily for clinical signs of distress, including significant weight loss, labored breathing, or lethargy. Humane endpoints were applied as necessary to minimize suffering.

### Fungal strains and media

*C. neoformans* strains KN99 and the clinical isolates UgCl422 and UgCl223(28) were used in this study. Fungal strains were cultured in 50 mL of yeast extract-peptone-dextrose (YPD) broth (1% yeast extract, 2% Bacto-peptone, and 2% dextrose) at 30°C with shaking at 300 rpm for 48 h. Following incubation, fungal cells were collected by centrifugation, washed with sterile, endotoxin-free phosphate-buffered saline (PBS), and enumerated using a hemocytometer.

### Mice

C57BL/6 (000664), BALB/c (000651), CBA/J (000656), Rag1^-/-^ (002216), and *Ccr2^-/^*^-^ (004999) mice were obtained from Jackson Laboratory (Bar Harbor, ME). Mice were 6 to 8 weeks old at the time of inoculation, and all experiments were conducted with an equal mix of both male and female mice to avoid sex bias.

### Neutrophil Depletions

Each mouse received intraperitoneal injections of 200 µg anti-Ly6G (clone 1A8), 100 µg RecombiMAb anti-Ly6G (1A8-CP129), or the corresponding isotype control antibody (Rat IgG2a (2A3) or RecombiMAb mouse IgG2a) all from Bio X Cell (Lebanon, NH) in 100 µL sterile PBS, beginning 24 h before infection and continuing every 48 h for the duration of the experiment. Depletion efficiency was assessed by flow cytometry in the lung, blood, and spleen 48 h after a single dose of antibody. The 1A8-CP129 antibody was used for all experiments other than those shown in Figure 1.

### Pulmonary inoculations

Mice were anesthetized with isoflurane and inoculated by orotracheal aspiration with 10^4^ or 10^5^ colony-forming units (CFU) of *C. neoformans* KN99, or 10^5^ CFU of the clinical isolates UgCI223 or UgCI422 in 50 μl of sterile PBS. Animals were monitored daily for clinical signs of disease. For survival studies, mice were euthanized when body weight fell below 80% of the weight recorded at the time of inoculation or when they became moribund. For fungal burden, cytokine, and flow cytometry studies, mice were euthanized by CO2 inhalation at the indicated times post-infection, together with uninfected (unchallenged) controls that received antibody but no fungus.

### Cytokine and chemokine analysis

Pulmonary cytokine and chemokine concentrations were quantified using a multiplex bead-based immunoassay. Lung tissue was excised and homogenized in 2 mL of ice-cold PBS containing 1× Pierce Protease Inhibitor cocktail (Thermo Scientific, Rockford, IL). Triton X-100 was then added to a final concentration of 0.05% and the samples were clarified by centrifugation. Supernatant fractions of the pulmonary homogenates were assayed with the ProcartaPlex Mouse Cytokine & Chemokine Panel 1, 26-plex according to the manufacturer’s instructions and acquired on a MAGPIX instrument. Concentrations were interpolated from standard curves run on each plate; values falling below the lower limit of quantification were assigned the limit value and are identified in the figures.

### Flow Cytometry

Pulmonary leukocyte populations were characterized by multiparameter flow cytometry. Briefly, lungs were harvested from individual mice and enzymatically digested for 30 min at 37°C in Dulbecco’s modified Eagle medium (DMEM) containing 0.05 mg/mL Liberase TM and 0.02 mg/mL DNase I (Roche, Indianapolis, IN). Following digestion, tissue suspensions were sequentially passed through sterile 70-μm and 40-μm nylon cell strainers (BD Biosciences, San Jose, CA) to obtain single-cell suspensions. Erythrocytes were lysed by incubating cell suspensions in ammonium chloride lysis buffer (0.859% NH₄Cl, 0.1% KHCO₃, and 0.0372% Na₂EDTA; pH 7.4; Sigma-Aldrich) for 3 min on ice. Lysis was terminated by adding a twofold excess of PBS. Cells were collected by centrifugation, resuspended in PBS, and stained with Fixable Viability Dye eFluor 506 (1:500; Invitrogen, Carlsbad, CA) for 30 min at 4°C in the dark. Following viability staining, cells were washed and resuspended in fluorescence-activated cell sorting (FACS) buffer (PBS containing 0.1% bovine serum albumin, 0.02% sodium azide, and 2 mM EDTA). Fc receptors were blocked by incubation with anti-CD16/CD32 (Fc Block; BD Biosciences, San Jose, CA) for 5 min to minimize nonspecific antibody binding. Cells at 1x10^6^ were incubated for 30 min at 4° C in the dark with optimal concentrations of fluorochrome-conjugated antibodies. The antibodies used for immunophenotyping, including their respective targets, clones, and manufacturers, are listed in Table S1. Following staining, cells were washed three times with FACS buffer and fixed in 2% ultrapure paraformaldehyde. Samples were acquired on a NovoCyte 3000 flow cytometer (Agilent Technologies, Santa Clara, CA), with more than 200,000 events collected per sample. Flow cytometry data were analyzed using FlowJo software (version 10; Tree Star, Ashland, OR). The gating strategies are shown in Figure S3. The absolute number of cells in each leukocyte subset was calculated by multiplying the absolute number of CD45+ cells by the percentage of cells stained for each population.

### Statistics

Data were analyzed with GraphPad Prism v10 (GraphPad Software, La Jolla, CA). Kaplan-Meier survival curves were compared by the Mantel-Cox log-rank test. Two-group comparisons at a given time point were made by t-test. Comparisons among more than two groups were performed using one-way analysis of variance (ANOVA), followed by Tukey’s multiple-comparisons test. The 26-plex cytokine panel was treated as a screen, and p-values were corrected across all analytes and time points within an experiment by the Benjamini-Hochberg false discovery rate procedure, and these are reported as q-values. Analytes for which 25% or more of the values in a group fell at the assay lower limit of quantification are flagged in the figures and were not used to support conclusions. A *P* value of less than 0.05 was considered statistically significant.

## ACKNOWLEDGMENTS

This work was supported by a National Institutes of Health grant K22 AI148724, the Center of Pediatric Experimental Therapeutics, and start-up funds from the Department of Clinical Pharmacy and Translational Science. The funders had no role in study design, data collection, interpretation, or the decision to submit the work for publication. We thank Dr. Kirsten Nielsen and Dr. David Meya for generously providing the *C. neoformans* clinical isolates UgCl422 and UgCl223.

